# Applying Cell Painting in Non-Tumorigenic Breast Cells to Understand Impacts of Common Chemical Exposures

**DOI:** 10.1101/2024.04.30.591893

**Authors:** Anagha Tapaswi, Nicholas Cemalovic, Katelyn M Polemi, Jonathan Z Sexton, Justin A Colacino

## Abstract

There are a substantial number of chemicals to which individuals in the general population are exposed which have putative, but still poorly understood, links to breast cancer. Cell Painting is a high-content imaging-based *in vitro* assay that allows for rapid and unbiased measurements of the concentration-dependent effects of chemical exposures on cellular morphology. We optimized the Cell Painting assay and measured the effect of exposure to 16 human exposure relevant chemicals, along with 21 small molecules with known mechanisms of action, for 48 hours in non-tumorigenic mammary epithelial cells, the MCF10A cell line. Through unbiased imaging analyses using CellProfiler, we quantified 3042 morphological features across approximately 1.2 million cells. We used benchmark concentration modeling to quantify significance and dose-dependent directionality to identify morphological features conserved across chemicals and find features that differentiate the effects of toxicants from one another. Benchmark concentrations were compared to chemical exposure biomarker concentration measurements from the National Health and Nutrition Examination Survey to assess which chemicals induce morphological alterations at human-relevant concentrations. Morphometric fingerprint analysis revealed similar phenotypes between small molecules and prioritized NHANES-toxicants guiding further investigation. A comparison of feature fingerprints via hypergeometric analysis revealed significant feature overlaps between chemicals when stratified by compartment and stain. One such example was the similarities between a metabolite of the organochlorine pesticide DDT (p,p’-DDE) and an activator of canonical Wnt signaling CHIR99201. As CHIR99201 is a known Wnt pathway activator and its role in ꞵ-catenin translocation is well studied, we studied the translocation of ꞵ-catenin following p’-p’ DDE exposure in an orthogonal high-content imaging assay. Consistent with activation of Wnt signaling, low dose p’,p’-DDE (25nM) significantly enhances the nuclear translocation of ꞵ-catenin. Overall, these findings highlight the ability of Cell Painting to enhance mode-of-action studies for toxicants which are common exposures in our environment but have previously been incompletely characterized with respect to breast cancer risk.

## INTRODUCTION

Since 2007, the rate of breast cancer-associated deaths has remained steady in women younger than 50, with a 2.6% lifetime mortality risk. The average risk for developing breast cancer is approximately 12% in the United States (Nicoletta et al. 2022). Breast cancer is the most common type of cancer in women worldwide, with the highest mortality rate has been observed in triple-negative breast cancer (TNBC) (Singh et al. 2021). TNBC is a particularly aggressive subtype of breast cancer that is negative for estrogen receptor (ER), progesterone receptor (PR), and does not overexpress human epidermal growth factor receptor 2 (HER2) (Howard et al. 2022). There are also substantial disparities in breast cancer incidence and outcomes by race/ethnicity in the US, with African American women being more likely to be diagnosed with triple-nega t ive breast cancers and more likely to die of breast cancer compared to women of other races/ethnicit ies (Newman et al. 2017). High-penetrance genetic risk factors are only thought to explain approximately 5-10% of aggressive breast cancer risk (Rodgers et al. 2018), thus there is a pressing need to identify additional modifiable risk factors that promote the development of these diseases.

A growing body of research points to environmental chemical exposures linked to an increased risk of breast cancer, including metals (cadmium and lead), perfluoroalkyl substances and pesticides (Koval et al. 2022). Developing new strategies to test chemical exposures, and their effects at population-relevant concentrations would provide important new methods for the identification of putative risk factors and their potential mechanisms of action.

Previous work in our lab, using data from the National Health and Nutrition Examina t ion Survey (NHANES), an ongoing population-based health study conducted by the US Centers for Disease Control and Prevention, identified a set of chemical exposures that are very commonly detected in US women, with additional stark differences in chemical exposure biomarker concentrations across women of various racial/ethnic groups (Nguyen et al. 2020). These chemicals (referred to as NHANES chemicals, here on) had concentrations significantly higher, on average, in African American women and included pesticide metabolites (2,5-dichlorophenol, 1,4-dichlorobenzene), chemicals of personal care products (methyl paraben, propyl paraben, monoethyl phthalate), and heavy metals (mercury and lead), which based on additional analysis of ToxCast high-throughput screening data, may be linked to breast cancer associated biological alterations (Polemi et al. 2021). Taking into consideration the common nature of exposure to these chemicals, we wanted to further investigate the effect of these on non-tumorigenic MCF10A breast cells in a high-throughput manner to quantify the concentration-dependent effects.

Morphological profiling describes the process of using automated imaging to quantify large feature sets, typically hundreds to thousands, from each experimental sample in a relatively unbiased way (Bray et al. 2016). High-Content Imaging (HCI) is a method of phenotypic screening for studying cells and sub-cellular components by simultaneously assessing multiple phenotypes in cell populations. Morphological HCI is customizable and can yield many measurements on a single cell level. Most large-scale imaging experiments extract only one or two features of cells, and/or aim to identify just a few ’hits’ in a screen, meaning that vast quantities of quantitative data about cellular state remain untapped (Bray et al. 2016). Bioactivity profiling has revealed that compounds that share mechanisms of action will induce similar phenotypic alterations in a cell (Svenningsen et al. 2019). The next-generation blueprint of computational toxicology at the U.S. Environmental Protection Agency (EPA) advocates for the use of non-targeted, high throughput profiling assays for the initial characterization of biological activity of environmental chemica ls (Willis et al. 2020). An image-based high throughput phenotypic profiling (HTPP) method, Cell Painting (Bray et al. 2016), is effective in quantifying subtle perturbations in cellular and subcellular features which can be implicated in disease onset and development and can be used to identify potential modes of action for chemical stressors (Willis et al. 2020). The Cell Painting assay is currently being deployed to novel human-derived cell types grown in monolayer in a straightforward manner, with minimal (or no) adjustment to sample preparation protocols when optimized across cell lines or laboratories (Willis et al. 2020). The method utilizes the power of automated fluorescence microscopy to measure 8 cellular components using six fluorescent stains which are imaged in 5 channels. The cellular components include DNA, RNA, nucleoli, endoplasmic reticulum, mitochondria, actin cytoskeleton, golgi apparatus, and the plasma membrane (Gustafsdottir et al. 2013). Bioactivity screening of environmental chemicals requires an approach that can measure the vast array of unknown phenotypes these chemicals may produce to detect and quantify biological effects across concentrations (Nyffeler et al. 2023).

In this study we aimed to adapt and establish, in our laboratory, Cell Painting as a morphometric study tool to investigate the concentration-responsive effects of a panel of 35 chemicals on MCF10A cells. We aimed to study the morphological effects of these chemicals and uncover potential commonalities between small molecules and NHANES chemicals of intere st. As the mode of action (MoAs) of these small molecules have been explored, similarities in the morphological perturbations between these two groups of chemicals could point towards potential shared MoAs for chemicals with undetermined mechanisms. We validate one such morphological similarity using an orthogonal high-content imaging assay to assess Wnt pathway activity. Finally, through concentration-response analyses, we compare the concentrations at which there are significant morphological perturbations with concentrations measured in the US population as part of NHANES.

## Materials and Methods

An overview of the experimental workflow is shown in Figure 1. Chemicals were purchased from Sigma-Aldrich (Saint Louis, MO, USA), Chem Service (West Chester, PA, USA) and Cayman Chemicals (Ann Arbor, MI, USA) (Table S1). 16% methanol free Paraformaldehyde was obtained from Fisher Scientific (Hampton, NH, USA). Hoechst 33342, Phalloidin (Alexa Fluor Plus 750), Concanavalin A (Alexa Fluor 594) and SYTO 14 Green were purchased from ThermoFisher (Waltham, MA, USA). Wheat Germ Agglutinin (CF®770) was purchased from Biotium (Hayward, CA, USA). Tissue culture treated black, clear bottom 384 well plates were purchased from Fisher Scientific (Hampton, NH, USA).

**Figure 1:**
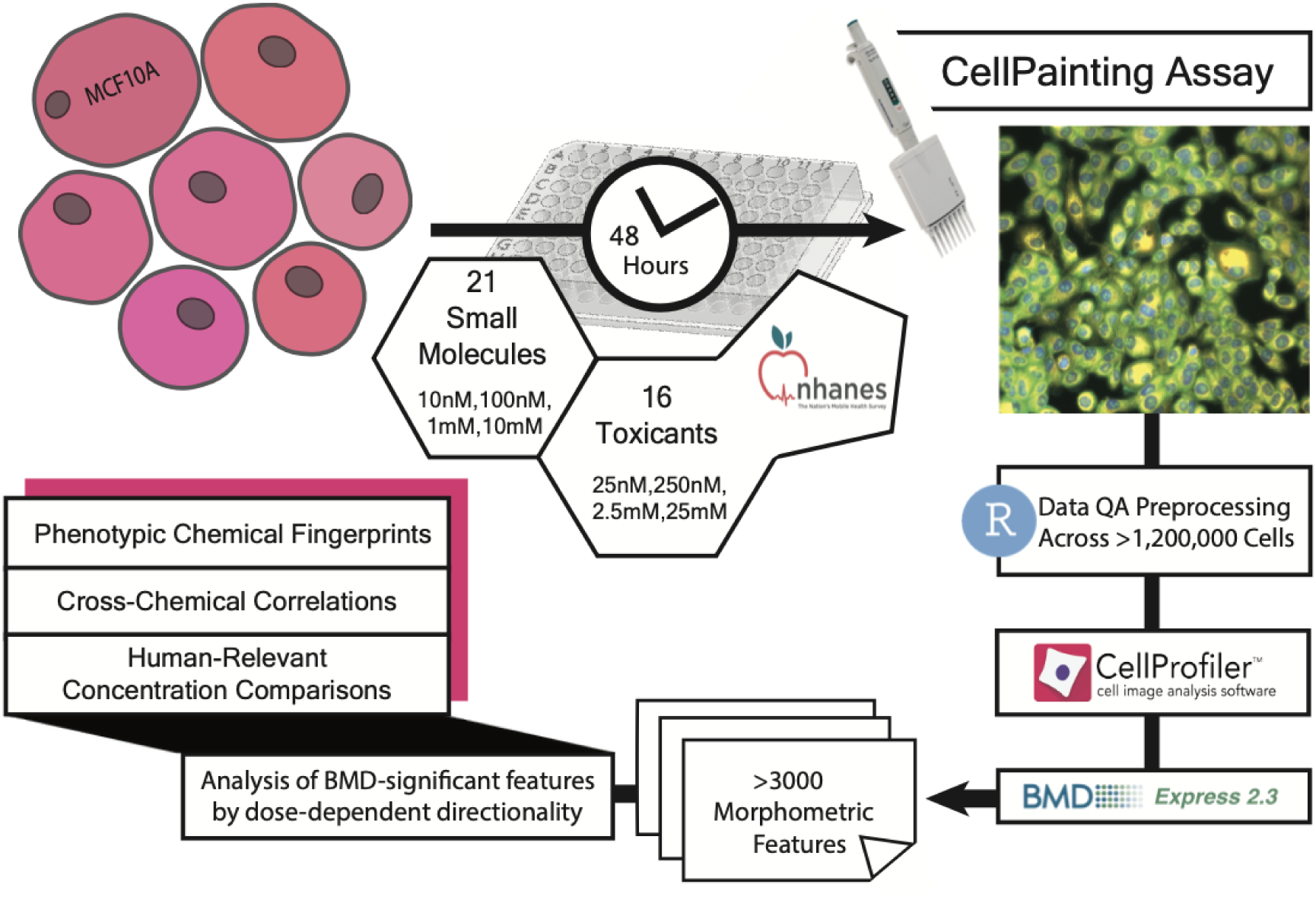
Methodology. A pictorial representation of the experimental and data analysis methods.

### Cell Culture and Plating

The non-malignant mammary epithelial cell line, MCF10A, was obtained from the American Type Culture Collection (ATCC, Manassas, VA, USA). MCF10A ce **l**s were cultured and maintained in Dulbecco’s Modified Eagle’s Medium/Ham’s F12 50/50 mix (Corning, Corning, NY, USA) supplemented with 5% Horse Serum (Thermo Fisher, Waltham, MA, USA), 2.5mg/mL HEPES (Thermo Fisher, Waltham, MA, USA), 5ug/mL Insulin (Gibco/Thermo Fisher, Waltham, MA, USA), 96ug/mL Hydrocortisone (StemCell, Vancouver, Canada), 100ug/mL Cholera toxin (Sigma-Aldrich, Saint Louis, MO, USA) and 20ng/mL Epidermal Growth Factor (StemCell, Vancouver, Canada). The cells were maintained at 37°C in a humidified 5% CO_2_ condition.

MCF10A cells were cultured and expanded at passage 105 for all rounds of cell culture in T-75 flasks (Corning, Corning, NY, USA). The cells were allowed to complete one passage in the flask prior to plating in black, clear flat bottom 384 well plates. 10uL of media was added to pre-warmed 384 well plates using a 2.5-125uL multi-channel pipette (Thermo Fisher Scientific, Waltham, MA). 500 cells/30 µL of media were seeded to each well at a slow speed to make up a final volume of 40 µL. The cells were left undisturbed in the cell culture hood for 1 hour to allow uniform distribution around the well and cells were allowed to attach in a 37°C humidified incubator with 5% CO_2_ for 24 hours prior to dosing. Cells were then dosed with the chemicals and incubated for 48 hours with 6 biological replicates per dose, along with appropriate vehicle controls.

### Compound Preparation and Dosing

The 35 chemicals under investigation were purchased from Sigma-Aldrich, Chem Service, and Cayman Chemicals (Table S1). The chemicals were weighed on a balance (Mettler Toledo, Columbus, OH, USA) at 5mg and dissolved in Dimethyl sulfoxide (DMSO) or in DNA/RNA free ultrapure Water (Invitrogen, Waltham, MA, USA) at a concentration of 5mg/mL and stored at -20°C for long term storage (Table S1). For dosing, an intermediate stock concentration of 5mM was prepared for NHANES toxicants and 2mM for small molecules, which were further diluted into complete MCF10A media, to final concentrations for dosing: 25uM, 2.5uM, 0.25uM and 0.025uM for NHANES chemicals and 10uM, 1uM, 0.1uM and 0.01uM for small molecules. These final concentrations were prepared fresh in DMEM/F12 complete media for each experiment. The final concentration of DMSO and water controls were at 0.5%, as were the prepared test concentrations.

### Cell Painting: Staining and Image Acquisition

The Cell Painting protocol was adapted from the Carpenter lab (Bray et.al, 2016) for use with our imaging platform, The Celllnsight CX5 High Content Screening Platform (Thermo Fisher Scientific, Waltham, MA, USA). The specific reagents we used are shown in Table S2. The stains Concanavalin A/Alex Fluor™ 594 conjugate (200 µg/mL), Pha **l**oidin/Alexa fluor™ Plus 750 (1:1000), and Wheat Germ Agglutinin/ CF®770 (5µg/mL) were changed from the original protocol to adapt for the fluorescent spectra available on the CX5. Briefly, after 48 hours of chemical dosing, the cell culture media was removed and the cells were washed with 40uL of 1X HBSS. After 1 wash, 35µL of 1X HBSS was removed, and 40µL of Mitotracker Deep Red (0.5µL/mL) prepared in DMEM/F12 complete media was added. Cells were incubated for 30 minutes in a 37°C humidified incubator with 5% CO_2_. Next, the cells were fixed using 4% methanol-free Paraformaldehyde (Alfa Aesar, Haverhill, MA, USA) for 20 minutes. The cells were washed once using a pre-set washing protocol on ELx405 Select CW (Biotek, Winooski, VT, USA) and permeabilized with 0.1% Triton X-100 for 15 minutes. The cells were washed two times and stained with a solution containing Hoechst 33342, Syto-14 green fluorescent nucleic acid stain, Concanavalin A, Mitotracker Deep Red FM, Phalloid in Alexa Plus™ 750, and Wheat Germ Agglutinin CF®770 for 30 minutes. The plate was washed 3 times and 40µL of fresh 1X HBSS was added to each well.

The stained plates were imaged using the Celllnsight CX5. The imaging was set up on Thermo Scientific HCS Studio: Cellomics Scan Version 6.6.1. Automated imaging was configured at a 20X/0.45 NA LUCPlan FLN objective with a target exposure set at 25%. Exposure times were selected based on the control wells. Nine fields were imaged at a single Z-height based on image-based autofocus on the first channel stained with Hoechst 33342 (350/461).

### Image QC and Illumination correction

Images were exported from the Cellnsight CX5 High Content Screening Platform in .C01 format. The open-source software Cell Profiler 4.2.1 was used for imaging quality control, processing, and analysis on a Windows-based workstation with dual Intel Xeon Platinum 8173M processors and 192 GB of RAM.

Raw images were run through a pipeline to perform illumination correction for all the acquired channels. An average model image was saved in .npy format for each channel. The set of raw images was then run through a pipeline set to calculate the saturation and blurriness parameters. The results were analyzed in Cell Profiler Analyst version 3.0.4 to get the threshold values for the set quality parameters. The upper and lower thresholds were saved for use in further analysis.

A cell segmentation and feature extraction pipeline was set up in CellProfiler, adapted from the Carpenter lab (Bray et al. 2016). This pipeline automatically carried out illumination correction and flagged images that failed to pass QC thresholds. Primary object segmentation was carried out using the Hoechst 33342 stain for the nucleus. Secondary objects were identified using the Syto 14 stain for Nuclear and Cytoplasmic RNA by Otsu thresholding. To complete morphological analysis, 3 regions of interest (ROIs) were identified: Nucleus, Cells, and Cytoplasm. Multiple parameters were analyzed including Colocalization, Granularity, Object Intensity, Intensity distribution, Neighbors, Object Size, Object Shape, and Texture within all the ROIs, for all the acquired stains. The data was exported using the Export to Spreadsheet module and saved as .csv files. A metadata file was prepared for each plate processed.

### Data preprocessing

Data from CellProfiler were imported into R statistical software for additional normalization, processing, and analysis. Briefly, single cell measurements in CSV format were quantified across the nuclei, cytoplasm, and cell body and were imported and merged with the experimental metadata containing information on the treatment, dose, and replicates per well. Features that had no information for any cells were removed. The Costes colocaliza t ion features were also removed from downstream analysis as they were often missing in more than 50% of cells. After this, cells with missing features were dropped to generate a complete dataset for analysis. To identify cytotoxic concentrations of chemicals that resulted in a significa nt perturbation in terms of cell number and viability, we identified doses of chemicals in which the average cell counts were less than 25% of the control we **l**s as “cytotoxic” to exclude these doses from morphological fingerprinting. Cell counts per well were quantified and saved as a csv file. For each plate, we performed normalization of features to the relevant control treatment (DMSO or water) using the *normalize* function in the Cytominer package (Caicedo et al. 2017), using median absolute deviation through the “robustize” operation. We then transformed the normalized data using the *transform* function in Cytominer, prior to aggregating the Cell Painting morphological profiles from a single cell to a per well basis using the *aggregate* function in Cytominer with the “median” operation. To prepare the aggregated data for exporting to BMDExpress2, data from wells with cytotoxic concentrations of chemicals were excluded and then the aggregated per we **l** morphological “fingerprints” were written to a tab-delimited text file where each column is a well with metadata including the treatment, the concentration, and the replicate.

### BMDExpress2 Analysis

Using BMDExpress2 (Phillips et al. 2019), we calculated concentration-response curves for each Cell Painting feature, for each chemical. Features were first filtered via one-way ANOVA, where features that did not meet a 2-fold change and unadjusted p-value < 0.05 cutoff were excluded from downstream analyses. For features that did meet the ANOVA filtering criteria, Hill, power, and polynomial models were fit for each feature to identify the best model for concentration-response. The benchmark response type was standard deviation, with a benchmark response factor of 1.349. We set the maximum iterations of the model to 250, with a confidence level of 0.95. Power was restricted to greater than or equal to 1 for the power model. Best fit models were chosen for each dose-response relationship per feature using a nested Chi-square test to select for the best model followed by lowest Akaike Information Criterion (AIC). Hill models were flagged if their ‘k’ parameter was sma **l**er than the lowest positive concentration, and flagged hill models were included in the data.

To visualize the estimated benchmark concentrations following concentration-response modeling, we generated benchmark concentration (BMC) accumulation plots using the R package ggplot2, filtering out features with a benchmark concentration higher than the highest concentration tested: 25 uM for NHANES-prioritized toxicants and 10 µM for small molecules. This plot allows us to compare bioactivity across chemicals, where chemicals that induce changes in more features at lower concentrations are considered more bioactive.

### Generation of total feature fingerprint across all chemicals

When conducting benchmark concentration-response modeling, for the best-predicted model fit for a given feature, BMDExpress2 identifies the direction of change (UP/DOWN). We used these direction of change data to develop a morphological “fingerprint” for each chemical which describes which features change in which direction within the concentration range tested. Given the concentratio n- dependent directionality of significant features, morphometric features that demonstrated no concentration-dependent significance across any compounds screened were removed from feature fingerprint visualization. This yielded 3042 morphometric features across our panel of small molecules and NHANES-prioritized toxicants. Feature fingerprints were visualized by heatmap using heatmap.2 function from the R package gplots. Morphological fingerprints were then compared via hierarchical clustering to group chemicals based on fingerprint similarity.

### Comparison of feature fingerprints via hypergeometric analysis

For each chemical, the significant features determined by BMC analysis were stratified by compartment (Cells, Cytoplasm, Nuclei) and stain (DNA, RNA, ER, Mito, AGP), generated from the R package gplots. We conducted a hypergeometric analysis of conserved features between different chemicals to quantify similarities in feature fingerprints. Features identified for significant concentration-dependent directionality, both up-regulated and down-regulated, were considered “significant” when comparing features between chemicals. Significant features shared between select small molecules and toxicants of interest were compared with one-sided Fisher’s exact tests in R. To investigate compartmental and organelle enrichment, all of the significant features shared between two chemicals were stratified for analysis by stain, compartment, as well across as 3042 features, reporting the numerical breakdown and Fisher’s exact test p-values.

### Comparison of NHANES Exposure Levels to Benchmark Concentrations

We compared concentrations of the tested chemicals to human biomarker concentrations measured as part of the National Health and Nutrition Examination Survey following previously established methodology (Nguyen et al. 2020, Polemi et al. 2021). For this analysis, we used our curated NHANES dataset, which contains information on chemical biomarker concentrations of the 15 assessed chemicals in up to 57786 women recruited from 1999-2018 (Vy Kim Nguyen et al. 2023). Chemical biomarker concentrations in women in NHANES were converted to molarity units to compare to the benchmark concentrations found *in vitro*. Overlapping NHANES concentrations and BMD concentrations indicate that relevant biomarkers measured in US women In NHANES occur within the benchmark doses found in cell painting. The chemical names, NHANES biomarker names and chemical code names were summarized in Table S6.

### ꞵ-Catenin immunofluorescence (IF) Staining and Translocation Quantification

To validate the fingerprint overlap between p,p’-DDE and the Wnt activating molecule CHIR99021, we conducted an orthogonal assay to quantify the translocation of ꞵ-catenin to the nucleus, a measure of activated Wnt signaling (Hwang et al. 2022, Kim et al. 2019). MCF10A cells were plated in 384 we **l** plates for 24 hours prior to dosing with p’,p’-DDE and CHIR99021. We treated MCF10A ce **l**s with 25, 2.5, 0.25 and 0.025uM p’,p’-DDE or 10, 1, 0.1, 0.01 uM CHIR99021, both for 48 hours. Vehicle control cells were treated with 0.5% DMSO. Cells were washed with 40uL of PBS and 35uL of PBS was removed, to avoid loss of cells that may be loosely attached due to chemical treatment. The cells were fixed with 4% Paraformaldehyde for 20 minutes and washed with PBS. Fixed cells were permeabilized with 0.1% Triton-X for 15 minutes and washed with PBS. The cells were blocked with 1% Bovine Serum Albumin (BSA) for 1 hour at room temperature and incubated with an anti-ꞵ-catenin antibody (Catalog no: 71-270-0, Thermo Fisher Scientific, Waltham, MA, USA) overnight at 4C. The cells were washed 3 times with 1X PBST before adding anti-Rabbit secondary antibody (Catalog No: ab150080, Abcam, Waltham, MA, USA) for 1 hour. Cells were washed 3 times and then incubated with Hoechst for 30 minutes to stain the nucleus. Cells were washed 3 times with 40µL 1X PBST and fresh 1X PBST was added prior to imaging.

The stained plate was imaged using the Celllnsight CX5 using Thermo Scientific HCS Studio: Cellomics Scan Version 6.6.1. Exposure times were selected based on the control wells. Nine fields were imaged at a single Z offset based on image-based autofocus on the first channel stained with Hoechst 33342 (350/461). Images were exported from the Cellnsight CX5 High Content Screening Platform in the acquired .C01 format. Cell Profiler 4.2.1 was used for imaging quality control, processing, and analysis. First, images were run through Illumination correction and QC pipeline to eliminate any background variations and filter out blurry and saturated images with set QC thresholds. The images were run through a cell segmentation and feature extraction pipeline using Hoechst 33342 stain to identify primary objects. Secondary objects were identified using the ꞵ-Catenin stain by Otsu thresholding. For complete morphological analysis, 3 regions of interest (ROIs) were identified: Nucleus, Cells and Cytoplasm. Features with respect to size, shape and intensity were collected within a **l** the ROIs, for Hoechst 33342 and ꞵ-catenin stains. The data was exported using the Export to Spreadsheet module and saved as .csv files. A metadata file was prepared. Using R Studio, the Mean Intensity for ꞵ-catenin stain in the nuclear region was subsetted out. Mean Intensity was averaged across replicates by treatment and concentration. The change in mean intensity was visualized using boxplots from the R package ggplot2 and a p value was determined by comparing nuclear mean intensity for control against the tested concentrations using the Wilcoxon statistical test.

## RESULTS

### Cell Painting Methodology and Cell Counts Data

We adapted and successfully established the Cell Painting protocol, modifying 2 dyes from the original protocol to best meet our requirements. Figure 2 displays example images of the stains in a control condition as well as following treatment with the chemotherapeutic agent and topoisomerase II inhibitor etoposide. Cell counts per well were quantified in data preprocessing and was visualized as a heat map for small molecules and NHANES chemicals across all doses. (Figure S1). We observed increased cell numbers, relative to control, at low doses for NHANES, chemicals like p,p’–DDE, bisphenol A (BPA), and cadmium chloride.

**Figure 2:**
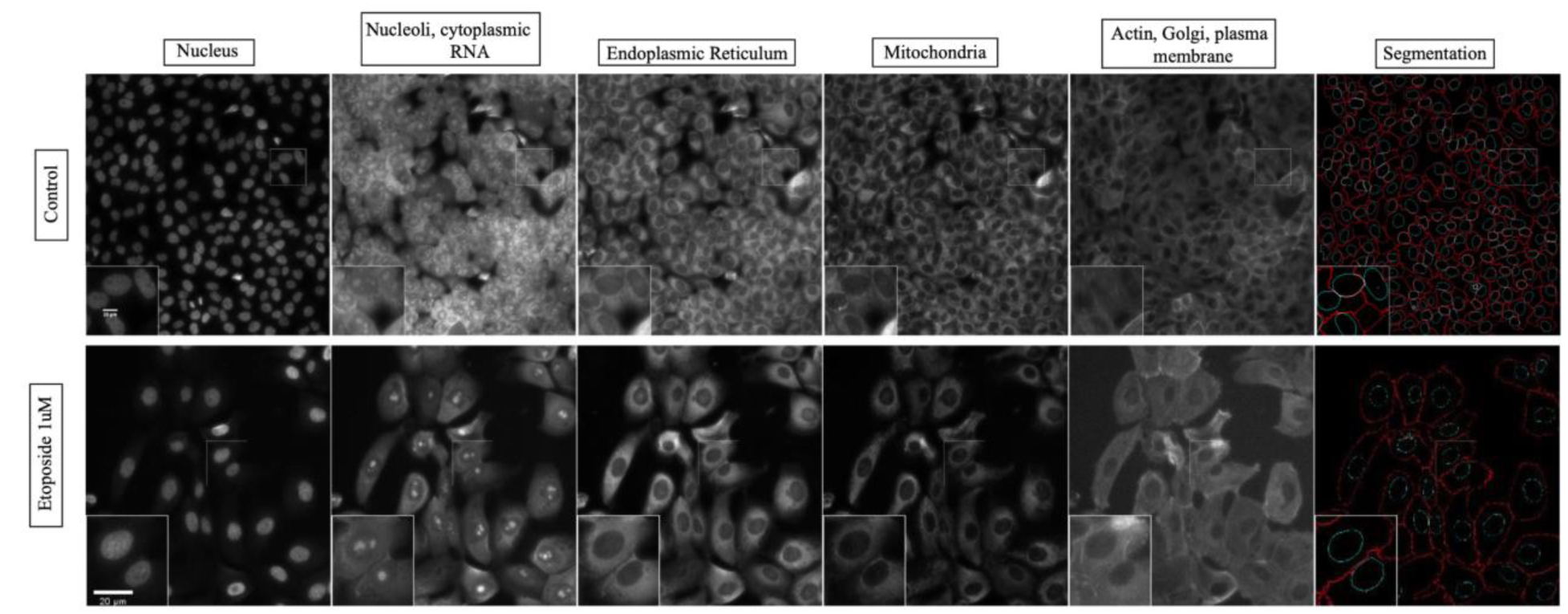
Cell Painting imaging panel in MCF10A cells. The columns show the 5 channels imaged in the Cell Painting protocol: Hoechst 33342 (DNA), Syto 14 (RNA), Concanavalin A (Endoplasmic Reticulum), Mitotracker (Mitochondria), Phalloidin (Actin) and WGA (Golgi and Plasma membrane). Supplementary table S2 outlines all the stains used in the experiment.

### Morphometric Fingerprint Analysis

Analysis of small molecules with validated targets, bioactivities, and phenotypes in tandem with NHANES-prioritized toxicants using CellPainting allows for rapid and unbiased quantification of altered morphological features and hypothesis generation of human-relevant effects of poorly-characterized toxicants. Clustering the 3042 morphometric features across the 35 compounds tested, the full feature fingerprint identified two distinct clusters of chemicals including a larger set of nuclear-active small molecules as well as a cluster of heavy metal toxicants and select pesticides (Figure 3A). This includes HDAC class I and II inhibitors like Vorinostat (Richon et al. 2006) and Trichostatin A (Kim M et al. 2019). The cluster also shows small molecules with known effects on mitochondrial function, namely, Menadione (Shandilya et al. 2021), FCCP (Grasmick et al. 2018) and Decitabine (Shin et al. 2012). Incorporating BMD-associated significance and concentration-dependent directionality allowed for the distinction of features that were consistently conserved across chemotypes and those features that distinguish toxicants from one another (Figure 3B). Feature and chemical clustering demonstrated Ce **l**Painting’s ability to characterize broad morphometric phenotypes of similar bioactivity as well as the sensitivity to distinguish morphometric fingerprints between structurally and functionally similar molecules. This is well observed within Figure 3A with the different histone deacetylase inhibitors (TSA, SAHA), nucleosides (5-Aza, Decitabine) clustering together. We observed fingerprint similarit ies between the metal/metalloid toxicants tested (sodium arsenite, mercury chloride, copper chloride, lead acetate).

**Figure 3:**
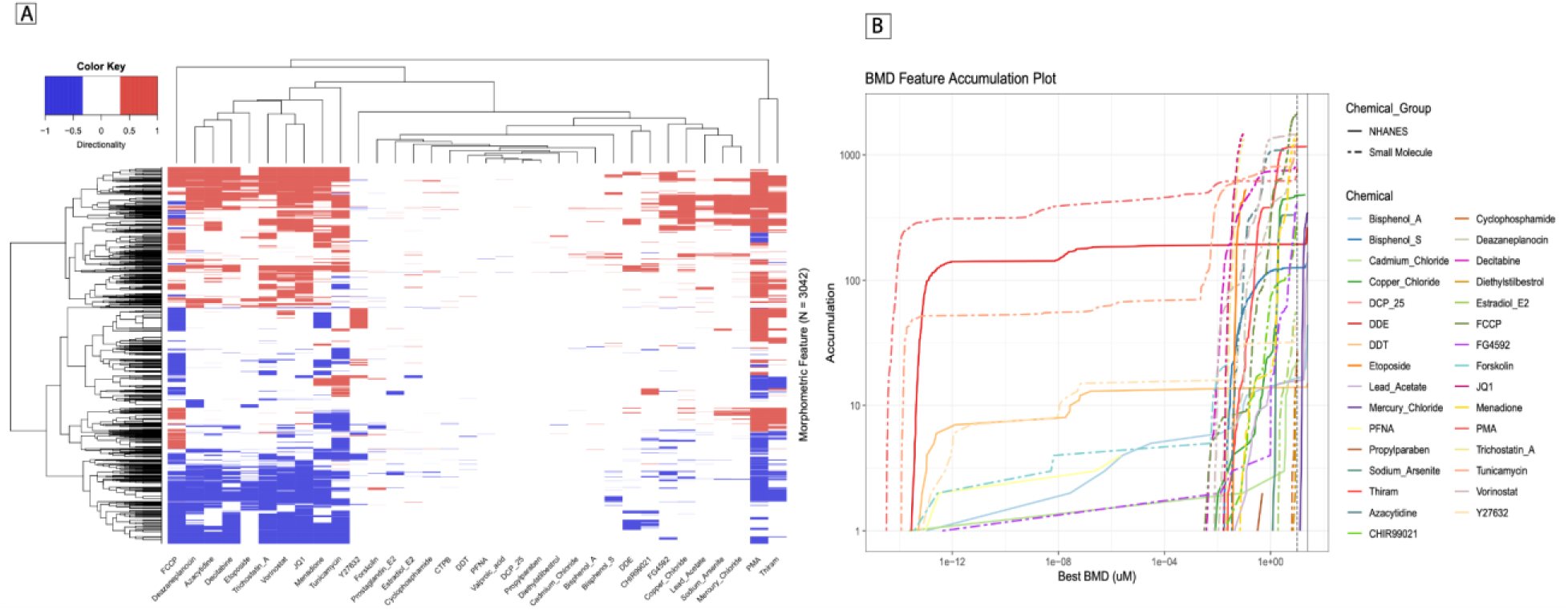
Morphological fingerprint analysis. (A) Heatmap depicting the morphological fingerprints for the small molecules and the NHANES chemicals, with fingerprints grouped using unbiased hierarchical clustering. (B) A benchmark concentration accumulation plot depicting the effects of tested chemicals, depicting the concentrations at which features are significantly changing relative to control for each assayed chemical.

Chemical clustering also revealed similar phenotypes between small molecules and NHANES- toxicants of interest, guiding further investigation. Chemical comparisons of interest were done to identify the total number of significant features depicting their up/down regulation. This analysis was done for the 3 ROIs (Table S3) and for the 5 individual stains (Table S4). These chemical comparisons were followed up with hypergeometric enrichment analysis of significant features across compartments and stains. This analysis allowed us to quantify the statistical significance of the overlap of dose-dependent features shared between two chemicals, typically a validated small molecule and an NHANES-prioritized toxicant (Table S5).

### Stain Enrichment Among Small Molecules and NHANES Prioritized Toxicants

Clustering within Figure 3A identified fingerprint similarity between the pesticide p,p’-DDE and the WNT activator CHIR99201. Of the 3042 morphometric features, p,p’-DDE and CHIR99201 shared 122 significant concentration-dependent features (p = 5.55E-63), 103 which resided in the nuclear compartment (p = 9.17E-81) and all 122 of which derived from the RNA stain channel (p = 7.52E-76) (Figure 4A). The morphometric feature fingerprint of FG-4592, a HIF prolyl hydroxylase inhibitor and HIF-1a stabilizer, showed the most similarity to fingerprints from heavy metal toxicants. As an example, copper chloride and FG-4592 shared 243 features (p = 7.31E- 114), 219 of which were from the mitochondrial channel (p = 1.54E-120) (Figure 4B). Additionally, PMA and Thiram shared 1070 features (p = 6.78E-246), 399 of which were associated with the endoplasmic reticulum channel (p = 4.80E-143) (Figure S2).

**Figure 4:**
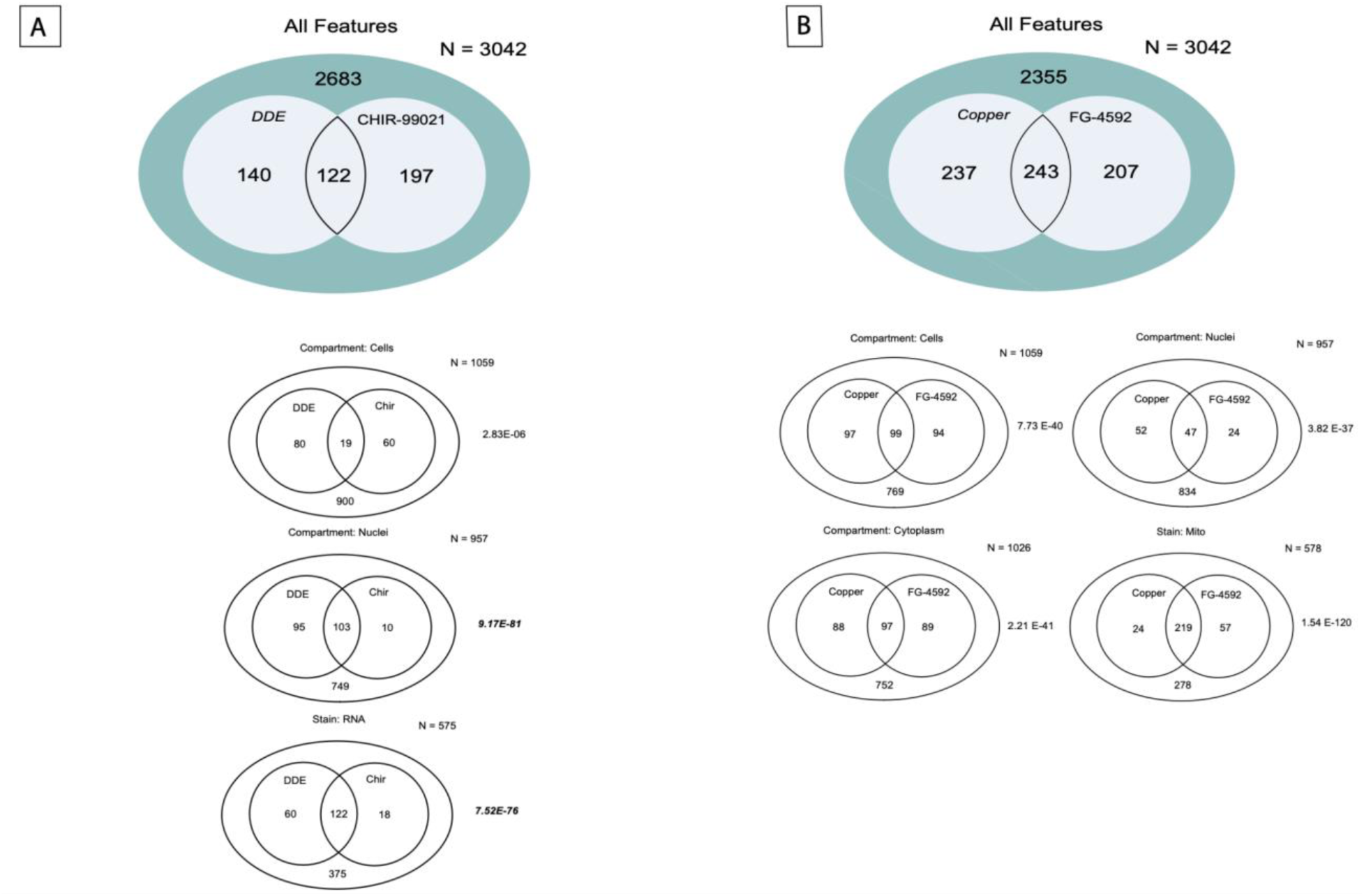
Chemical comparison Venn diagrams. 4A and 4B: Venn diagrams representing the overlap of common features between 2 chemicals combinations: DDE and CHIR99021 and Copper and FG-4592 with in different Regions of Interest: Cells, Nuclei and Cytoplasm along with the significant cellular stain showing maximum overlapping features. Supplementary table S3 and S4 bring together all the features that were significantly different in regions of interest: Nuclei, Cells and Cytoplasm in all the 5 tested stains. Supplementary table S5 presents the chemical combinations having a significant overlap in common features with the tested P values.

### β-Catenin Nuclear Translocation

Given the significant features shared between p,p’-DDE and CHIR99201, we next wanted to investigate whether there is a potential mode of action shared between the 2 chemicals. As CHIR99201 is a known WNT activator, we analyzed β-catenin expression and localization after treatment with the pesticide p,p’-DDE. Translocation of β-catenin from the cytoplasm to the nucleus is consistent with activation of the canonical Wnt pathway (Liu et al. 2022).

Post-treatment, we analyzed the ꞵ-catenin mean intensity in the nuclear region. Treatment with 25nM and 250nM p,p’-DDE for 48 hours significantly increased the ꞵ-catenin intensity relative to vehicle control (p=0.0097 and 0.0135, respectively, Figure 5A). Translocation of β-catenin was confirmed by a significant increase in intensity when cells were treated with CHIR99201 at 10uM (p=0.00011), as expected as the experimental positive control. The data showed translocation of ꞵ-catenin from cytoplasm to nucleus and decreased membrane expression as compared to control (Figure 5B).

**Figure 5:**
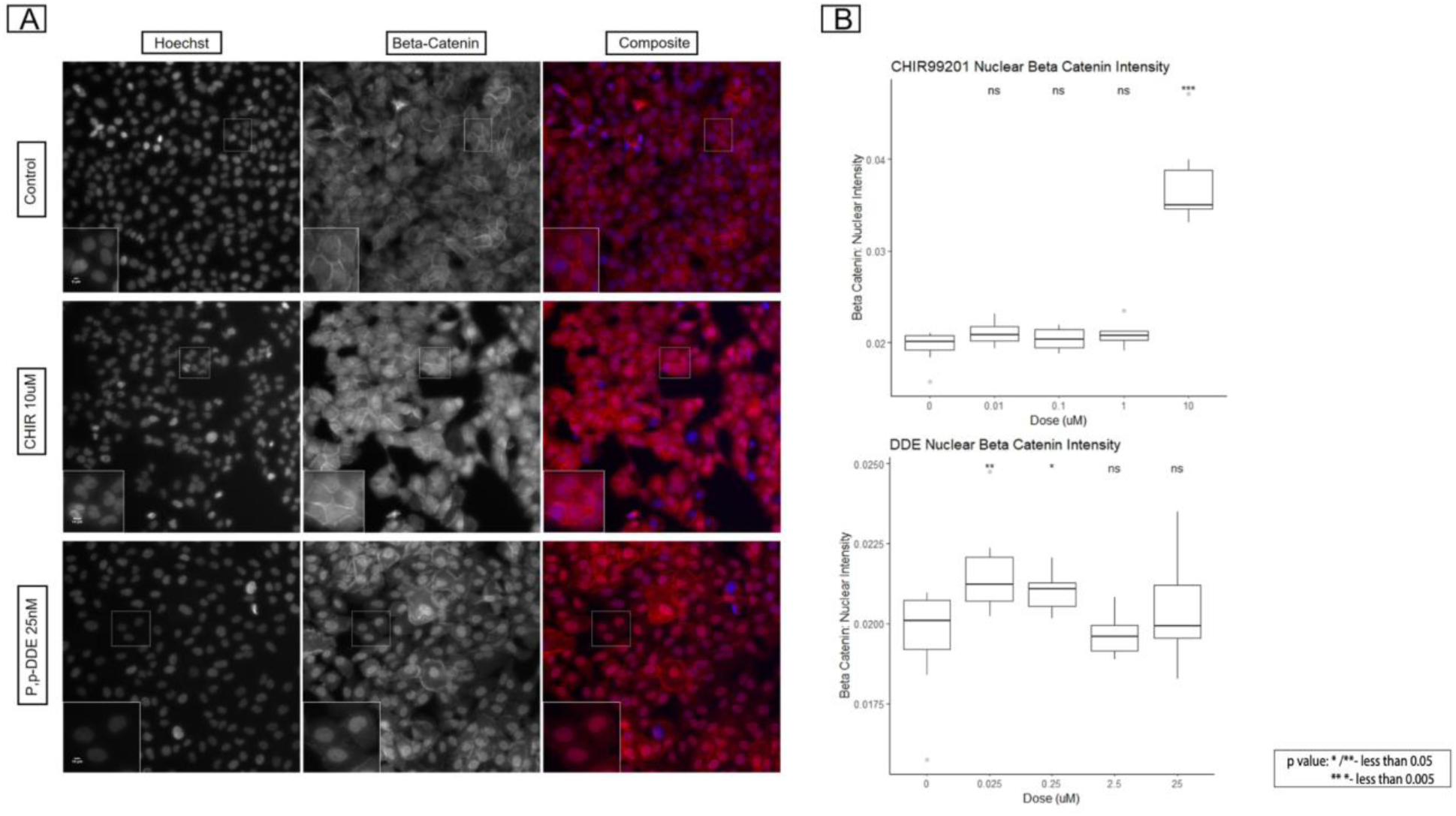
ꞵ-Catenin expression and observed translocation on treatment with low dose p,p’- DDE. 5A: Immunofluorescence images depicting the expression of ꞵ-Catenin in Control, CHIR99201 (10uM) and DDE (25nM). 5B: Analysis of ꞵ-Catenin intensity in the nuclear region in MCF10A cells. P values were determined using the Wilcoxon rank-sum test. *P<0.05, **P<0.01, ****P<0.0001

### Comparison Between Benchmark Concentrations and NHANES Chemical Biomarker Concentrations

We next compared the tested *in vitro* concentrations to exposure biomarker levels detected in individuals in the United States. We compared our benchmark dose concentrations from our in- vitro experiments to National Health and Nutrition Examination Survey (NHANES) biomarker concentration data for each chemical, plotting the range of blood or urine chemical biomarker concentrations in US individuals compared to the range of the BMCs for each chemical from our in-vitro data (Figure 6). The strongest overlaps between BMCs and NHANES were observed for p,p’-DDE, copper chloride, and sodium arsenite, although many other chemicals had morphological features that change at concentrations that are detectable in exposure biomarker concentrations detected in NHANES study participants.

**Figure 6.**
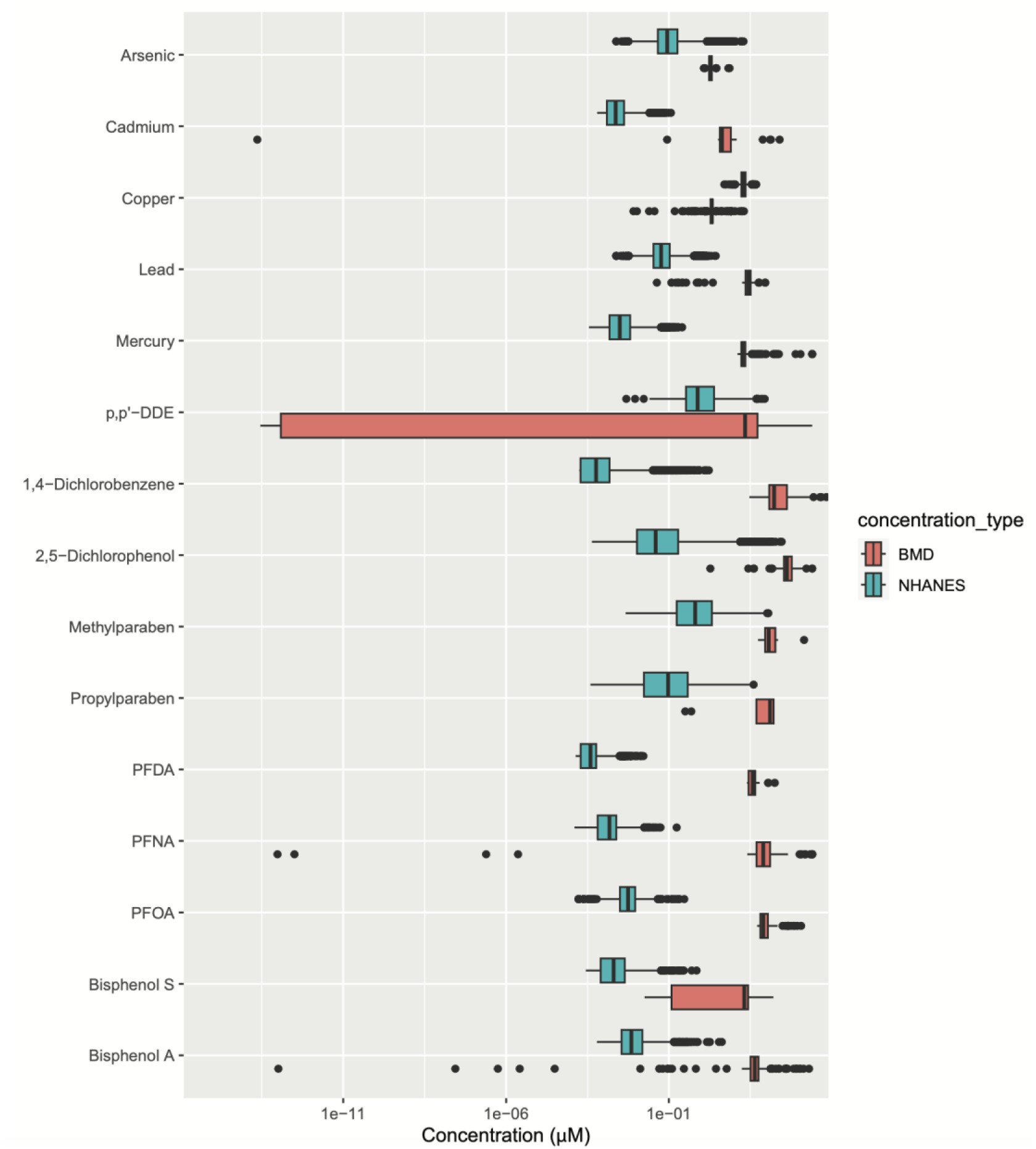
A comparative bar plot depicting chemical biomarker concentrations in women in NHANES were converted to molarity units to compare to the benchmark dose concentrations found in cell painting to understand the biologically-relevant exposure doses. Supplementary table S6 shows the NHANES biomarker and chemical codenames for the chemicals having biologically relevant exposure doses.

## DISCUSSION

There are many common chemical exposures that individuals can encounter on a daily basis with unclear links to breast cancer risk. Here, we adapted the Cell Painting assay, a morphological analysis tool to study the effect of toxicants that are commonly detected in US women and compared their effects to known small molecules in non-tumorigenic breast cells. Among the 35 chemicals tested, we observed meaningful small molecule-NHAN ES chemical morphologica l fingerprint combinations that share a significant overlap in altered features. Lower concentration effects for p,p’–DDE, sodium arsenite, and copper chloride fall within the range of the NHANES human-relevant concentrations. These findings highlight that biomarker concentrations of some chemicals that are commonly detected in US women are high enough to induce morphological alterations in non-tumorigenic breast cells *in vitro*.

Morphometric fingerprint analysis revealed significant small molecule-NHANES chemical combinations which show potential commonalities between their modes of action. Cell Painting provided us with an in-depth method of quantifying phenotypic differences across regions of interest (ROI) namely, in the nucleus, cell, and cytoplasm, and across 5 different stains, allowing visualization of 8 different organelles. The ROI-based phenotypic analysis highlights the strength of the cell painting as a tool to decipher unique and shared phenotypic features between chemical exposures. Modifications in different regions of interest can inform us of biological mechanisms of a given exposure. When put together, effects across different ROIs can contribute significant information regarding the potential MOAs for test chemicals.

We identified a statistically significant overlap in feature fingerprints between p,p’–DDE and CHIR99201. CHIR99021 is the most commonly used GSK-3b inhibitor and is considered the standard small molecule agonist of the Wnt pathway (Law et al. 2022). CHIR99021 is a potent and selective GSK-3α/β inhibitor and Wnt/β-catenin signaling pathway activator that stabilizes β- catenin and activates Wnt/β-catenin pathway (Naujok et al. 2014). Previous work in colorectal adenocarcinoma cells revealed that total and nuclear ꞵ-catenin was significantly increased following p,p’-DDE exposure (Song et al. 2014). Similarly here in MCF10A cells, we also observed an increased translocation of ꞵ-catenin from cytoplasm to nucleus. Concurrently, we also observed decreased expression of ꞵ-catenin in cell membranes in cells treated with lower doses of p,p’–DDE. These observations highlight that low dose p,p’–DDE may activate Wnt signaling, which may explain, in part, the increased cell numbers observed in low dose p,p’–DDE exposures. These findings may also provide a potential mechanistic link to support epidemiological findings of associations between early-life exposure to DDT and risk of breast cancer later in life, which would be worthy of further investigation (Cohn et al. 2015, Cohn et al. 2007).

A significant number of features were also shared between copper chloride and FG4592. FG-4592 is a known hypoxia-inducible factor prolyl hydroxylase inhibitor (HI-PHI) (Provenzano et al. 2016). The role of hypoxia and inhibition of EGLN1, a hypoxia-inducible factor prolyl hydroxylase, by FG-4592 has been described earlier in both pancreatic and ovarian cancer (Fujimoto et al. 2019, Price et al. 2019). The role of HIF-1A being inhibited by cadmium has been observed in human primary breast organoids, where we identified that HIF-1A was essential for mammary organoid formation (Rocco et al. 2018). Characterizing how heavy metals interact with hypoxia signaling will be an important area of future research to understand potential mechanis t ic links between breast development and breast cancer.

Thiram is a commonly used fungicide. Thiram’s effects have not been actively studied in breast cells previously, it has been studied in avian growth plate chondrocytes with respect to its effect on MMP-2 and MMP-9 (Simsa et al. 2007). Thiram, in addition to PMA, induced activity of MMP-9 in chicken chondrocytes resulting in the formation of a nonvascularized, non-mineralized plaque in the growth plate (Simsa et al. 2007). PMA induces MMP-9 expression via a protein kinase Cα(PKCα)-dependent signaling cascade in BEAS-2B human lung epithelial cells (Shin et al. 2007). Stimulation of MCF-7 cells with PMA significantly downregulated the expression of hsa-miR-204-5p and significantly upregulated the mRNA expression of human MMP-9 (Farhana et al. 2023). An interesting future direction would be to explore whether part of Thiram’s MOA involved upregulation of MMP-9 expression.

One major limitation of this study is that we assessed chemical effects at one exposure duration (48hrs) and in one cell line (MCF10A). This initial study, although successful in identifying dose-dependent morphological alterations, also opens avenues to further include more time points and cell lines from additional individuals to understand interindividual heterogeneity in response to common environmental stressors. This work is currently underway in our laboratory. Another limitation with reference to concentrations of chemical biomarkers measured in NHANES is that these biomarkers are studied in blood and urine, which could or could not be reflective of the levels in the mammary tissue. Additionally, this *in vitro* analysis of the test chemicals may not reflect what happens in the context of a tissue *in vivo*. Additional investigation of the effects of these chemicals in the context of the breast tissue microenvironment would provide complementary information about how these exposures may impact breast cell biology and cancer risk.

The emergence of multiple chemical combinations that show a significant number of shared features shines a light on the ability of Cell Painting to serve as a morphometric analysis tool in the context of toxicants with incompletely understood MOAs. Future goals for our work in this area include adapting Cell Painting and the subsequent data analysis methods in human-derived normal mammary cells from diverse donors, to better understand interindividual responses to environmental stressors. Overall, Cell Painting is a highly informative and cost-effective strategy for the concentration-dependent analysis of human exposure-relevant chemicals with poorly characterized MOAs.

## Acknowledgments

This work was supported by grants from the National Institutes of Health (R01 ES028802, T32 ES00706, P30 ES017885, R01 AG072396, P30 CA046592, S10 OD034245). The content is solely the responsibility of the authors and does not necessarily represent the official views of the National Institutes of Health.

## Author Contributions

**Anagha Tapaswi**: Conceptualization, Methodology, Validation, Formal analysis, Investigat io n, Data Curation, Writing-Original Draft, Visualization. **Nicholas Cemalovic**: Formal analysis, Data Curation, Writing-Original Draft, Visualization, Validation, Methodology. **Katelyn M. Polemi**: Formal analysis, Visualization, Validation, Methodology, Writing- Review and Editing. **Jonathan Z Sexton**: Writing- Review and Editing. **Justin A Colacino**: Conceptualization, Methodology, Formal Analysis, Data Curation, Validation, Visualization, Writing-Review and Editing, Supervision, Project administration and Funding acquisition.

