## Supplemental Tables and Figures for "Applying Cell Painting in Non-Tumorigenic Breast Cells to Understand Impacts of Common Chemical Exposures"

**Small Molecules:**

| Chemical | CAS Number | Vendor; Catalog number | Solvent |
|---|---|---|---|
| 5-Azacytidine | 320-67-2 | Cayman Chemicals; 11164 | Dimethyl Sulfoxide (DMSO) |
| Decitabine | 2353-33-5 | Cayman Chemicals; 11166 | Dimethyl Sulfoxide (DMSO) |
| Vorinostat | 149647-78-9 | Cayman Chemicals; 10009929 | Dimethyl Sulfoxide (DMSO) |
| Valproic Acid | 1069-66-5 | Cayman Chemicals; 13033 | Dimethyl Sulfoxide (DMSO) |
| Trichostatin A | 58880-19-6 | Cayman Chemicals; 89730 | Dimethyl Sulfoxide (DMSO) |
| CTPB | 586976-24-1 | Cayman Chemicals; 19570 | Dimethyl Sulfoxide (DMSO) |
| 3- Deazaneplanocin | 120964-45-6 | Cayman Chemicals; 11102 | Dimethyl Sulfoxide (DMSO) |
| Etoposide | 33419-42-0 | Cayman Chemicals; 11102 | Dimethyl Sulfoxide (DMSO) |
| Cyclophosphamide | 50-18-0 | Cayman Chemicals; 13849 | Dimethyl Sulfoxide (DMSO) |
| FCCP | 370-86-5 | Cayman Chemicals; 15218 | Dimethyl Sulfoxide (DMSO) |
| Diethylstilbestrol | 56-53-1 | Cayman Chemicals; 10006876 | Dimethyl Sulfoxide (DMSO) |
| Y-27632 | 146986-50-7 | Cayman Chemicals; 10005583 | Dimethyl Sulfoxide (DMSO) |
| CHIR99021 | 252917-06-9 | Cayman Chemicals; 13122 | Dimethyl Sulfoxide (DMSO) |
| Forskolin | 66575-29-9 | Cayman Chemicals; 11018 | Dimethyl Sulfoxide (DMSO) |
| Prostaglandin E2 | 363-24-6 | Cayman Chemicals; | Dimethyl Sulfoxide |

| | | | |
|---|---|---|---|
| | | 14010 | (DMSO) |
| Mendione | 58-27-5 | Cayman Chemicals; M5625 | Dimethyl Sulfoxide (DMSO) |
| Thapsigargin | 67526-95-8 | Cayman Chemicals; 10522 | Dimethyl Sulfoxide (DMSO) |
| Tunicamycin | 11089-65-9 | Cayman Chemicals; 11445 | Dimethyl Sulfoxide (DMSO) |
| PMA | 16561-29-8 | Cayman Chemicals; 10008014 | Dimethyl Sulfoxide (DMSO) |
| FG4592 | 808118-40-3 | Cayman Chemicals; 15294 | Dimethyl Sulfoxide (DMSO) |
| JQ1 (+) | 1268524-70-4 | Cayman Chemicals; 11187 | Dimethyl Sulfoxide (DMSO) |

**NHANES:**

| **Chemical** | **CAS Number** | **Vendor; Catalog number** | **Solvent** |
|---|---|---|---|
| Methylparaben | 99-76-3 | Sigma Aldrich; 47889 | Dimethyl Sulfoxide (DMSO) |
| Propylparaben | 94-13-3 | Sigma Aldrich; 1577008 | Dimethyl Sulfoxide (DMSO) |
| Thiram | 137-26-8 | Sigma Aldrich; 45689 | Dimethyl Sulfoxide (DMSO) |
| 2,5-Dichlorophenol | 583-78-8 | Sigma Aldrich; D70007 | Dimethyl Sulfoxide (DMSO) |
| 1,4-Dichlorobenzene | 99106-46-7 | Sigma Aldrich; D56829 | Dimethyl Sulfoxide (DMSO) |
| Sodium (meta) Arsenite | 7784-46-5 | Sigma Aldrich; S7400 | Water |
| Lead (II) Acetate Trihydrate | 6080-56-4 | Sigma Aldrich; 316512 | Water |

| | | | |
|---|---|---|---|
| Mercury (II) Chloride | 7487-94-7 | Sigma Aldrich; 215465 | Water |
| Copper (II) Chloride | 7447-39-4 | Sigma Aldrich; 222011 | Water |
| Cadmium Chloride | 10108-64-2 | Sigma Aldrich; 202908 | Water |
| Bisphenol S | 80-09-1 | Sigma Aldrich; 103039 | Dimethyl Sulfoxide (DMSO) |
| Bisphenol A | 9980-05-7 | Chem Service; S-F7295S | Dimethyl Sulfoxide (DMSO) |
| PFDA | 335-76-2 | Sigma Aldrich; 177741 | Dimethyl Sulfoxide (DMSO) |
| PFNA | 375-95-1 | Sigma Aldrich; 394459 | Dimethyl Sulfoxide (DMSO) |
| DDE | 72-55-9 | Chem Service; N-10875 | Dimethyl Sulfoxide (DMSO) |
| DDT | 8017-34-3 | Chem Service; N-11567 | Dimethyl Sulfoxide (DMSO) |

**Supplemental Table S1: Chemical Information**: *Small molecules and NHANES chemicals chosen and their associated CAS number, catalog number, vendor and solvents.*

| **Cellular Stain** | **Filter (excitation; nm)** | **Filter (emission; nm)** | **Cellular component** | **Cell profiler channel name** |
|---|---|---|---|---|
| Hoechst 33342 | 350 | 461 | Nucleus | DNA |
| SYTO 14 Green Fluorescent Nucleic Acid stain | 521 | 547 | Nucleoli, Cytoplasmic RNA | RNA |
| Concanavalin A/Alex Fluor™ 594 conjugate | 590 | 617 | Endoplasmic reticulum (α-mannopyranosyl and α-glucopyranosyl residues) | ER |
| Mito Tracker Deep Red FM | 644 | 665 | Mitochondria | Mito |
| Phalloidin/ Alex Fluor™ Plus 750,<br>Wheat Germ Agglutinin/ CF®770 | 758/770 | 784/797 | F-Actin cytoskeleton, Golgi apparatus, Plasma membrane | AGP |

**Supplemental Table S2: Stain Information**: *Cell Painting stains selected and their associated fluorophores, and stained cellular compartments.*

| Chemical | Compartment | num_upreg | num_downreg | total_num_sig |
| --- | --- | --- | --- | --- |
| Azacytidine | Cells | 242 | 238 | 480 |
| Azacytidine | Cytoplasm | 176 | 232 | 408 |
| Azacytidine | Nuclei | 114 | 87 | 201 |
| Bisphenol_A | Cells | 11 | 3 | 14 |
| Bisphenol_A | Cytoplasm | 19 | 3 | 22 |
| Bisphenol_A | Nuclei | 8 | 0 | 8 |
| Bisphenol_S | Cells | 39 | 24 | 63 |
| Bisphenol_S | Cytoplasm | 35 | 39 | 74 |
| Bisphenol_S | Nuclei | 19 | 1 | 20 |
| Cadmium_Chloride | Cells | 6 | 3 | 9 |
| Cadmium_Chloride | Cytoplasm | 5 | 0 | 5 |
| Cadmium_Chloride | Nuclei | 9 | 0 | 9 |
| CHIR99021 | Cells | 57 | 22 | 79 |
| CHIR99021 | Cytoplasm | 33 | 10 | 43 |
| CHIR99021 | Nuclei | 93 | 104 | 197 |
| Copper_Chloride | Cells | 163 | 33 | 196 |
| Copper_Chloride | Cytoplasm | 157 | 28 | 185 |
| Copper_Chloride | Nuclei | 88 | 11 | 99 |
| CTPB | Cells | 5 | 3 | 8 |
| CTPB | Cytoplasm | 6 | 7 | 13 |
| CTPB | Nuclei | 4 | 4 | 8 |
| Cyclophosphamide | Cells | 8 | 12 | 20 |
| Cyclophosphamide | Cytoplasm | 9 | 7 | 16 |
| Cyclophosphamide | Nuclei | 12 | 3 | 15 |
| DCB_14 | Cells | 0 | 0 | 0 |
| DCB_14 | Cytoplasm | 0 | 0 | 0 |
| DCB_14 | Nuclei | 0 | 0 | 0 |
| DCP_25 | Cells | 0 | 0 | 0 |
| DCP_25 | Cytoplasm | 1 | 0 | 1 |

| | | | | |
|---|---|---|---|---|
| DCP_25 | Nuclei | 0 | 0 | 0 |
| DDE | Cells | 63 | 36 | 99 |
| DDE | Cytoplasm | 49 | 1 | 50 |
| DDE | Nuclei | 36 | 77 | 113 |
| DDT | Cells | 1 | 5 | 6 |
| DDT | Cytoplasm | 1 | 3 | 4 |
| DDT | Nuclei | 2 | 3 | 5 |
| Deazaneplanocin | Cells | 193 | 256 | 449 |
| Deazaneplanocin | Cytoplasm | 153 | 240 | 393 |
| Deazaneplanocin | Nuclei | 153 | 187 | 340 |
| Decitabine | Cells | 155 | 225 | 380 |
| Decitabine | Cytoplasm | 147 | 187 | 334 |
| Decitabine | Nuclei | 67 | 122 | 189 |
| Diethylstilbestrol | Cells | 0 | 2 | 2 |
| Diethylstilbestrol | Cytoplasm | 13 | 3 | 16 |
| Diethylstilbestrol | Nuclei | 1 | 0 | 1 |
| Estradiol_E2 | Cells | 11 | 17 | 28 |
| Estradiol_E2 | Cytoplasm | 22 | 15 | 37 |
| Estradiol_E2 | Nuclei | 6 | 3 | 9 |
| Etoposide | Cells | 79 | 114 | 193 |
| Etoposide | Cytoplasm | 79 | 117 | 196 |
| Etoposide | Nuclei | 61 | 78 | 139 |
| FCCP | Cells | 296 | 449 | 745 |
| FCCP | Cytoplasm | 284 | 452 | 736 |
| FCCP | Nuclei | 242 | 385 | 627 |
| FG4592 | Cells | 124 | 69 | 193 |
| FG4592 | Cytoplasm | 124 | 62 | 186 |
| FG4592 | Nuclei | 60 | 11 | 71 |
| Forskolin | Cells | 13 | 21 | 34 |
| Forskolin | Cytoplasm | 28 | 10 | 38 |

| | | | | |
|---|---|---|---|---|
| Forskolin | Nuclei | 5 | 24 | 29 |
| JQ1 | Cells | 337 | 318 | 655 |
| JQ1 | Cytoplasm | 307 | 265 | 572 |
| JQ1 | Nuclei | 130 | 148 | 278 |
| Lead_Acetate | Cells | 49 | 2 | 51 |
| Lead_Acetate | Cytoplasm | 97 | 2 | 99 |
| Lead_Acetate | Nuclei | 13 | 4 | 17 |
| Menadione | Cells | 231 | 326 | 557 |
| Menadione | Cytoplasm | 250 | 326 | 576 |
| Menadione | Nuclei | 150 | 292 | 442 |
| Mercury_Chloride | Cells | 122 | 5 | 127 |
| Mercury_Chloride | Cytoplasm | 113 | 5 | 118 |
| Mercury_Chloride | Nuclei | 99 | 1 | 100 |
| Methylparaben | Cells | 0 | 0 | 0 |
| Methylparaben | Cytoplasm | 0 | 0 | 0 |
| Methylparaben | Nuclei | 0 | 0 | 0 |
| PFDA | Cells | 0 | 0 | 0 |
| PFDA | Cytoplasm | 0 | 0 | 0 |
| PFDA | Nuclei | 0 | 0 | 0 |
| PFNA | Cells | 0 | 2 | 2 |
| PFNA | Cytoplasm | 0 | 2 | 2 |
| PFNA | Nuclei | 0 | 0 | 0 |
| PMA | Cells | 488 | 398 | 886 |
| PMA | Cytoplasm | 466 | 336 | 802 |
| PMA | Nuclei | 369 | 176 | 545 |
| Propylparaben | Cells | 0 | 0 | 0 |
| Propylparaben | Cytoplasm | 2 | 0 | 2 |
| Propylparaben | Nuclei | 0 | 0 | 0 |
| Prostaglandin_E2 | Cells | 4 | 27 | 31 |
| Prostaglandin_E2 | Cytoplasm | 3 | 42 | 45 |

| | | | | |
|---|---|---|---|---|
| Prostaglandin_E2 | Nuclei | 3 | 2 | 5 |
| Sodium_Arsenite | Cells | 112 | 4 | 116 |
| Sodium_Arsenite | Cytoplasm | 99 | 2 | 101 |
| Sodium_Arsenite | Nuclei | 113 | 2 | 115 |
| Thiram | Cells | 287 | 143 | 430 |
| Thiram | Cytoplasm | 239 | 131 | 370 |
| Thiram | Nuclei | 269 | 94 | 363 |
| Trichostatin_A | Cells | 235 | 291 | 526 |
| Trichostatin_A | Cytoplasm | 229 | 272 | 501 |
| Trichostatin_A | Nuclei | 150 | 195 | 345 |
| Tunicamycin | Cells | 254 | 335 | 589 |
| Tunicamycin | Cytoplasm | 264 | 335 | 599 |
| Tunicamycin | Nuclei | 235 | 305 | 540 |
| Valproic_acid | Cells | 0 | 0 | 0 |
| Valproic_acid | Cytoplasm | 0 | 0 | 0 |
| Valproic_acid | Nuclei | 1 | 2 | 3 |
| Vorinostat | Cells | 347 | 287 | 634 |
| Vorinostat | Cytoplasm | 276 | 269 | 545 |
| Vorinostat | Nuclei | 128 | 160 | 288 |
| Y27632 | Cells | 89 | 34 | 123 |
| Y27632 | Cytoplasm | 98 | 43 | 141 |
| Y27632 | Nuclei | 52 | 40 | 92 |

**Supplemental Table S3: Chemical Comparisons in Regions of Interest (ROIs)**: *Significant features up /downregulated in Nucleus, Cells and Cytoplasm.*

| | | | | |
|---|---|---|---|---|
| Azacytidine | AGP | 18 | 37 | 55 |
| Azacytidine | DNA | 106 | 221 | 327 |
| Azacytidine | ER | 62 | 55 | 117 |
| Azacytidine | Mito | 163 | 12 | 175 |
| Azacytidine | RNA | 122 | 164 | 286 |
| Bisphenol_A | AGP | 5 | 0 | 5 |
| Bisphenol_A | DNA | 0 | 0 | 0 |
| Bisphenol_A | ER | 20 | 0 | 20 |
| Bisphenol_A | Mito | 9 | 0 | 9 |
| Bisphenol_A | RNA | 4 | 0 | 4 |
| Bisphenol_S | AGP | 2 | 0 | 2 |
| Bisphenol_S | DNA | 37 | 54 | 91 |
| Bisphenol_S | ER | 15 | 0 | 15 |
| Bisphenol_S | Mito | 10 | 0 | 10 |
| Bisphenol_S | RNA | 18 | 5 | 23 |
| Cadmium_Chloride | AGP | 2 | 0 | 2 |
| Cadmium_Chloride | DNA | 0 | 0 | 0 |
| Cadmium_Chloride | ER | 11 | 0 | 11 |
| Cadmium_Chloride | Mito | 0 | 0 | 0 |
| Cadmium_Chloride | RNA | 6 | 2 | 8 |
| CHIR99021 | AGP | 5 | 0 | 5 |
| CHIR99021 | DNA | 6 | 0 | 6 |
| CHIR99021 | ER | 1 | 7 | 8 |
| CHIR99021 | Mito | 101 | 22 | 123 |
| CHIR99021 | RNA | 49 | 91 | 140 |
| Copper_Chloride | AGP | 20 | 0 | 20 |
| Copper_Chloride | DNA | 7 | 0 | 7 |
| Copper_Chloride | ER | 97 | 8 | 105 |
| Copper_Chloride | Mito | 217 | 26 | 243 |
| Copper_Chloride | RNA | 45 | 7 | 52 |
| CTPB | AGP | 0 | 0 | 0 |
| CTPB | DNA | 1 | 0 | 1 |

| | | | | |
|---|---|---|---|---|
| CTPB | ER | 0 | 7 | 7 |
| CTPB | Mito | 0 | 0 | 0 |
| CTPB | RNA | 4 | 0 | 4 |
| Cyclophosphamide | AGP | 3 | 0 | 3 |
| Cyclophosphamide | DNA | 1 | 0 | 1 |
| Cyclophosphamide | ER | 3 | 3 | 6 |
| Cyclophosphamide | Mito | 0 | 8 | 8 |
| Cyclophosphamide | RNA | 12 | 4 | 16 |
| DCB_14 | AGP | 0 | 0 | 0 |
| DCB_14 | DNA | 0 | 0 | 0 |
| DCB_14 | ER | 0 | 0 | 0 |
| DCB_14 | Mito | 0 | 0 | 0 |
| DCB_14 | RNA | 0 | 0 | 0 |
| DCP_25 | AGP | 1 | 0 | 1 |
| DCP_25 | DNA | 0 | 0 | 0 |
| DCP_25 | ER | 0 | 0 | 0 |
| DCP_25 | Mito | 0 | 0 | 0 |
| DCP_25 | RNA | 0 | 0 | 0 |
| DDE | AGP | 0 | 0 | 0 |
| DDE | DNA | 0 | 0 | 0 |
| DDE | ER | 70 | 0 | 70 |
| DDE | Mito | 0 | 0 | 0 |
| DDE | RNA | 75 | 107 | 182 |
| DDT | AGP | 3 | 7 | 10 |
| DDT | DNA | 0 | 0 | 0 |
| DDT | ER | 0 | 0 | 0 |
| DDT | Mito | 0 | 0 | 0 |
| DDT | RNA | 0 | 0 | 0 |
| Deazaneplanocin | AGP | 15 | 16 | 31 |
| Deazaneplanocin | DNA | 121 | 301 | 422 |
| Deazaneplanocin | ER | 20 | 52 | 72 |

| | | | | |
|---|---|---|---|---|
| Deazaneplanocin | Mito | 137 | 16 | 153 |
| Deazaneplanocin | RNA | 136 | 232 | 368 |
| Decitabine | AGP | 13 | 0 | 13 |
| Decitabine | DNA | 83 | 137 | 220 |
| Decitabine | ER | 22 | 43 | 65 |
| Decitabine | Mito | 77 | 9 | 86 |
| Decitabine | RNA | 125 | 285 | 410 |
| Diethylstilbestrol | AGP | 14 | 4 | 18 |
| Diethylstilbestrol | DNA | 0 | 1 | 1 |
| Diethylstilbestrol | ER | 0 | 0 | 0 |
| Diethylstilbestrol | Mito | 0 | 0 | 0 |
| Diethylstilbestrol | RNA | 0 | 0 | 0 |
| Estradiol_E2 | AGP | 0 | 0 | 0 |
| Estradiol_E2 | DNA | 8 | 4 | 12 |
| Estradiol_E2 | ER | 15 | 25 | 40 |
| Estradiol_E2 | Mito | 0 | 0 | 0 |
| Estradiol_E2 | RNA | 12 | 6 | 18 |
| Etoposide | AGP | 12 | 27 | 39 |
| Etoposide | DNA | 112 | 254 | 366 |
| Etoposide | ER | 10 | 0 | 10 |
| Etoposide | Mito | 52 | 1 | 53 |
| Etoposide | RNA | 1 | 4 | 5 |
| FCCP | AGP | 164 | 340 | 504 |
| FCCP | DNA | 113 | 261 | 374 |
| FCCP | ER | 230 | 63 | 293 |
| FCCP | Mito | 133 | 221 | 354 |
| FCCP | RNA | 132 | 341 | 473 |
| FG4592 | AGP | 0 | 0 | 0 |
| FG4592 | DNA | 31 | 12 | 43 |
| FG4592 | ER | 9 | 0 | 9 |
| FG4592 | Mito | 222 | 54 | 276 |

| | | | | |
|---|---|---|---|---|
| FG4592 | RNA | 40 | 45 | 85 |
| Forskolin | AGP | 6 | 0 | 6 |
| Forskolin | DNA | 23 | 4 | 27 |
| Forskolin | ER | 9 | 41 | 50 |
| Forskolin | Mito | 0 | 0 | 0 |
| Forskolin | RNA | 0 | 0 | 0 |
| JQ1 | AGP | 14 | 6 | 20 |
| JQ1 | DNA | 111 | 240 | 351 |
| JQ1 | ER | 203 | 58 | 261 |
| JQ1 | Mito | 248 | 58 | 306 |
| JQ1 | RNA | 126 | 292 | 418 |
| Lead_Acetate | AGP | 21 | 6 | 27 |
| Lead_Acetate | DNA | 0 | 0 | 0 |
| Lead_Acetate | ER | 63 | 0 | 63 |
| Lead_Acetate | Mito | 64 | 0 | 64 |
| Lead_Acetate | RNA | 11 | 0 | 11 |
| Menadione | AGP | 116 | 287 | 403 |
| Menadione | DNA | 100 | 209 | 309 |
| Menadione | ER | 89 | 157 | 246 |
| Menadione | Mito | 140 | 9 | 149 |
| Menadione | RNA | 124 | 213 | 337 |
| Mercury_Chloride | AGP | 19 | 1 | 20 |
| Mercury_Chloride | DNA | 0 | 0 | 0 |
| Mercury_Chloride | ER | 139 | 1 | 140 |
| Mercury_Chloride | Mito | 149 | 2 | 151 |
| Mercury_Chloride | RNA | 26 | 0 | 26 |
| Methylparaben | AGP | 0 | 0 | 0 |
| Methylparaben | DNA | 0 | 0 | 0 |
| Methylparaben | ER | 0 | 0 | 0 |
| Methylparaben | Mito | 0 | 0 | 0 |
| Methylparaben | RNA | 0 | 0 | 0 |

| | | | | |
|---|---|---|---|---|
| PFDA | AGP | 0 | 0 | 0 |
| PFDA | DNA | 0 | 0 | 0 |
| PFDA | ER | 0 | 0 | 0 |
| PFDA | Mito | 0 | 0 | 0 |
| PFDA | RNA | 0 | 0 | 0 |
| PFNA | AGP | 0 | 2 | 2 |
| PFNA | DNA | 0 | 0 | 0 |
| PFNA | ER | 0 | 0 | 0 |
| PFNA | Mito | 0 | 0 | 0 |
| PFNA | RNA | 0 | 0 | 0 |
| PMA | AGP | 325 | 166 | 491 |
| PMA | DNA | 87 | 193 | 280 |
| PMA | ER | 375 | 148 | 523 |
| PMA | Mito | 363 | 170 | 533 |
| PMA | RNA | 105 | 148 | 253 |
| Propylparaben | AGP | 0 | 0 | 0 |
| Propylparaben | DNA | 2 | 0 | 2 |
| Propylparaben | ER | 0 | 0 | 0 |
| Propylparaben | Mito | 0 | 0 | 0 |
| Propylparaben | RNA | 0 | 0 | 0 |
| Prostaglandin_E2 | AGP | 2 | 0 | 2 |
| Prostaglandin_E2 | DNA | 0 | 7 | 7 |
| Prostaglandin_E2 | ER | 0 | 0 | 0 |
| Prostaglandin_E2 | Mito | 5 | 40 | 45 |
| Prostaglandin_E2 | RNA | 2 | 22 | 24 |
| Sodium_Arsenite | AGP | 17 | 0 | 17 |
| Sodium_Arsenite | DNA | 0 | 0 | 0 |
| Sodium_Arsenite | ER | 116 | 1 | 117 |
| Sodium_Arsenite | Mito | 170 | 1 | 171 |
| Sodium_Arsenite | RNA | 18 | 0 | 18 |
| Thiram | AGP | 130 | 37 | 167 |
| Thiram | DNA | 89 | 113 | 202 |

| | | | | |
|---|---|---|---|---|
| Thiram | ER | 312 | 100 | 412 |
| Thiram | Mito | 205 | 41 | 246 |
| Thiram | RNA | 24 | 24 | 48 |
| Trichostatin_A | AGP | 79 | 128 | 207 |
| Trichostatin_A | DNA | 121 | 245 | 366 |
| Trichostatin_A | ER | 178 | 31 | 209 |
| Trichostatin_A | Mito | 48 | 24 | 72 |
| Trichostatin_A | RNA | 121 | 263 | 384 |
| Tunicamycin | AGP | 76 | 95 | 171 |
| Tunicamycin | DNA | 53 | 114 | 167 |
| Tunicamycin | ER | 112 | 228 | 340 |
| Tunicamycin | Mito | 326 | 119 | 445 |
| Tunicamycin | RNA | 135 | 347 | 482 |
| Valproic_acid | AGP | 0 | 2 | 2 |
| Valproic_acid | DNA | 0 | 0 | 0 |
| Valproic_acid | ER | 0 | 0 | 0 |
| Valproic_acid | Mito | 1 | 0 | 1 |
| Valproic_acid | RNA | 0 | 0 | 0 |
| Vorinostat | AGP | 84 | 80 | 164 |
| Vorinostat | DNA | 84 | 185 | 269 |
| Vorinostat | ER | 132 | 57 | 189 |
| Vorinostat | Mito | 260 | 28 | 288 |
| Vorinostat | RNA | 122 | 282 | 404 |
| Y27632 | AGP | 208 | 93 | 301 |
| Y27632 | DNA | 0 | 0 | 0 |
| Y27632 | ER | 24 | 12 | 36 |
| Y27632 | Mito | 0 | 7 | 7 |
| Y27632 | RNA | 0 | 0 | 0 |

**Supplemental Table S4: Chemical Comparisons by individual stains**: *Significant features up/downregulated in DNA, RNA, ER, Mito and AGP*.

| Chemical Comparison | | N = 3042 total features | Feature Compartment (N = 3042) | | | Feature Stain (N = 2,811*) | | | | |
|---|---|---|---|---|---|---|---|---|---|---|
| **Chemical 1 (N = # significant figures)** | **Chemical 2 (N = # significant figures)** | **# of shared significant features (one-sided ficher's test p-value)** | **Cells N = 1059 total features** | **Cytoplasm N = 1026 total features** | **Nuclei N = 957 total features** | **DNA N = 521 total features** | **RNA N = 575 total features** | **Mito N = 578 total features** | **ER N = 568 total features** | **AGP N = 569 total features** |
| DDE (N = 262) | CHIR99021 (N = 293) | 122 (5.55E-63) | 19 (2.83 E-06) | 0 | 103 (9.17E-81) | 0 | 122 (7.52E-76) | 0 | 0 | 0 |
| Thiram (N = 1163) | PMA (N = 2233) | 1070 (6.78E-246) | 415 (2.06 E-95) | 356 (1.41E-85) | 299 (2.86E-81) | 181 (1.87 E-73) | 45 (3.31E-17) | 215 (1.18E-21) | 399 (4.80E-143) | 157 (5.55E-22) |
| Copper_Chloride (N = 480) | FG4592 (N = 450) | 243 (7.31E-114) | 99 (7.73 E-40) | 97 (2.21E-41) | 47 (3.82E-37) | 0 | 10 (0.06) | 219 (1.54E-120) | 4 (0.013) | 0 |
| Sodium_Arsenite (N = 332) | Mercury_Chloride (N = 345) | 293 (3.28E-321) | 108 (7.97 E-125) | 97 (2.08E-116) | 88 (6.75E-93) | 0 | 18 (<E-300) | 143 (7.49E-116) | 113 (8.70E-100) | 15 (1.09E-26) |
| Copper_Chloride (N = 480) | Mercury_Chloride (N = 345) | 275 (6.70E-207) | 104 (5.87 E-88) | 106 (2.71E-88) | 65 (4.30E-54) | 0 | 21 (4.01E-22) | 142 (1.01E-73) | 91 (1.61E-64) | 18 (1.52E-33) |
| Copper_Chloride (N = 480) | Sodium_Arsenite (N = 332) | 258 (4.39E-186) | 103 (4.62 E-82) | 96 (3.19E-85) | 59 (2.89E-39) | 0 | 17 (4.21E-21) | 141 (1.60E-56) | 80 (2.04E-54) | 17 (<E-300) |

**Supplemental Table S5: Overlap of dose dependent features**: *Statistical significance of the overlap of dose-dependent features shared between two chemicals, typically a validated small molecule and a NHANES prioritized chemical.*

| **Chemical Name** | **NHANES Biomarker** | **Chemical Codename** |
|---|---|---|
| Lead (II) Acetate Trihydrate | Blood lead (ug/dL) | LBXBPB |
| Copper (II) Chloride | Serum Copper (ug/dL) | LBXSCU |
| Sodium (meta) Arsenite | Urinary arsenic, total (ug/L) | URXUAS |
| Mercury (II) Chloride | Blood mercury, total (ug/L) | LBXTHG |
| Cadmium Chloride | Blood cadmium (ug/L) | LBXBCD |
| Bisphenol A | Urinary Bisphenol A (ng/mL) | URXBPH |
| Bisphenol S | Urinary Bisphenol S (ug/L) | URXBPS |
| Methylparaben | Methyl paraben (ng/ml) | URXMPB |
| Propylparaben | Propyl paraben (µg/L) | URXPPB |
| PFDA | Perfluorodecanoic acid | LBXPFDE |
| PFNA | Perfluorononanoic acid | LBXPFNA |
| Thiram | Urinary 2-Thioxothiazolidine-4-carboxylic acid (ng/mL) | URXTTC |
| 2,5-Dichlorophenol | Urinary 2,5-dichlorophenol (µg/L) | URX14D |
| 1,4-Dichlorobenzene | Blood 1,4-Dichlorobenzene (ng/mL) | LBXVDB |
| DDE | ppDDE Lipid Adjusted (ng/g) | LBXPDELA |
| DDT | ppDDT lipid Adj (ng/g) | LBXPDELA |

**Supplemental Table S6: NHANES Biomarker Information:** *Chemical used in-vitro and their corresponding exposure biomarker codenames in NHANES.*

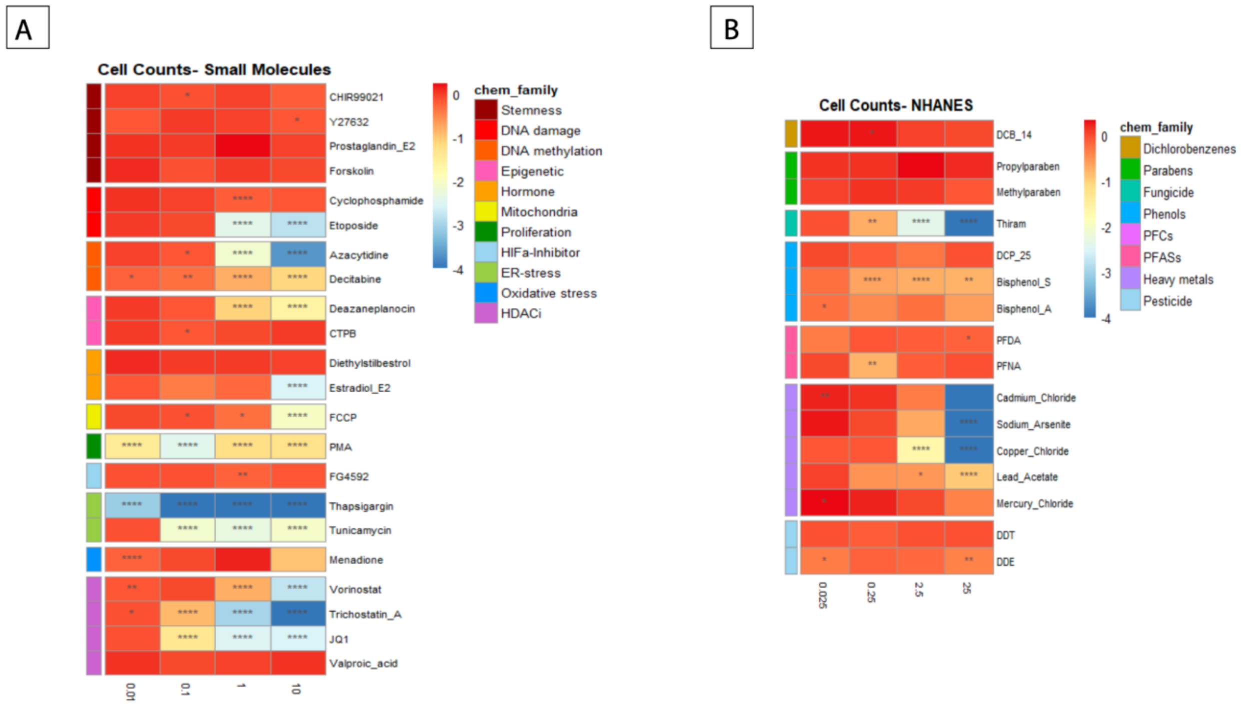


**Supplementary Figure S1: Cell Counts**: *A heatmap representation of the cell counts normalized by control cell counts for the tested small molecules and NHANES chemicals. P values were determined using the Wilcoxon test. *P<0.05, **P<0.01, ****P<0.0001*

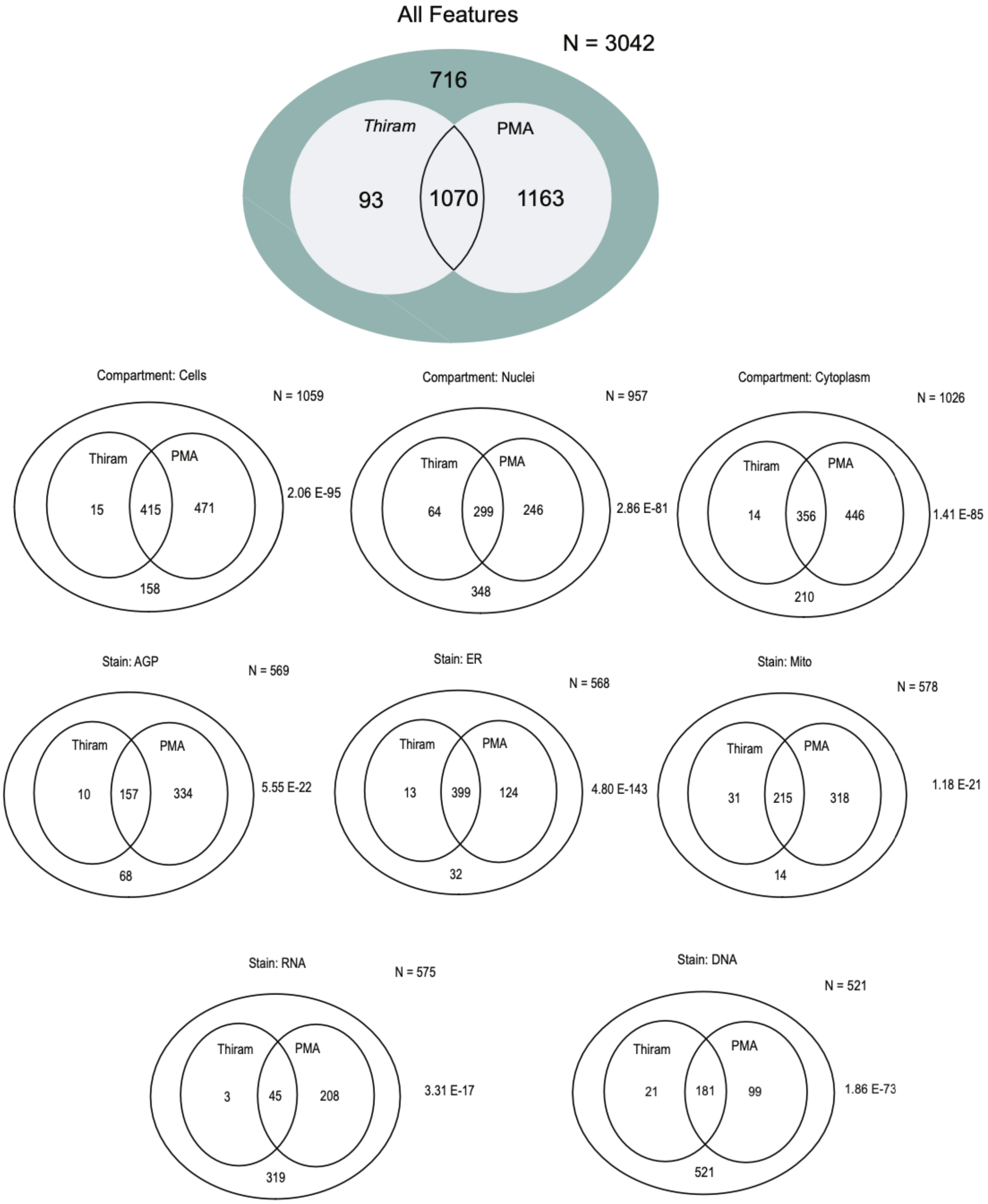


**Supplementary Figure S2: Overlap between Thiram and PMA:** *A Venn diagram representing the overlap of common features between 2 chemical combinations: PMA and Thiram within different Regions of Interest: Cells, Nuclei and Cytoplasm along with the significant cellular stain showing maximum overlapping features.*
